# Genetic removal of p70 S6K1 corrects coding sequence length-dependent alterations in mRNA translation in fragile X syndrome mice

**DOI:** 10.1101/2020.04.26.062281

**Authors:** Sameer Aryal, Francesco Longo, Eric Klann

**Author notes:** Correspondence to: Eric Klann.

## Abstract

Loss of the fragile X mental retardation protein (FMRP) causes fragile X syndrome (FXS). FMRP is widely thought to repress protein synthesis, but its translational targets and modes of control remain in dispute. We previously showed that genetic removal of p70 S6 kinase 1 (S6K1) corrects altered protein synthesis as well as synaptic and behavioral phenotypes in FXS mice. In this study, we examined the gene-specificity of altered mRNA translation in FXS and the mechanism of rescue with genetic reduction of S6K1 by carrying out ribosome profiling and RNA-Seq on cortical lysates from wild-type, FXS, S6K1 knockout, and double knockout mice. We observed reduced ribosome footprint abundance in the majority of differentially translated genes in the cortices of FXS mice. We used molecular assays to discover evidence that the reduction in ribosome footprint abundance reflects an increased rate of ribosome translocation, which is captured as a decrease in the number of translating ribosomes at steady state, and is normalized by inhibition of S6K1. We also found that genetic removal of S6K1 prevented a positive-to-negative gradation of alterations in translation efficiencies (RF/mRNA) with coding sequence length across mRNAs in FXS mouse cortices. Our findings reveal the identities of dysregulated mRNAs and a molecular mechanism by which reduction of S6K1 prevents altered translation in FXS.

## Introduction

Loss of expression or function of the fragile X mental retardation protein (FMRP) causes fragile X syndrome (FXS), the most prevalent inherited form of intellectual disability and the leading monogenic cause of autism. FMRP is an RNA-binding protein whose primary function is widely believed to repress mRNA translation in neurons. Accordingly, elevated net *de novo* protein synthesis is observed in the brains of FXS model mice (Qin et al., 2005). Precise control of translation is especially critical in the brain because rapid protein synthesis underlies long-lasting synaptic plasticity, multiple forms of which are impaired in FXS mice (Sidorov et al., 2013). There is consensus that dysregulated translation underlies a majority of phenotypes exhibited by FXS mice, including autism-like behaviors (Darnell and Klann, 2013). Restoring translational homeostasis in the brain therefore presents an attractive therapeutic concept in FXS. Accordingly, genetic deletion of the translation-stimulator p70 S6 kinase 1 (S6K1) in FXS mice rescues multiple phenotypes, including exaggerated translation, aberrant synaptic plasticity and dendritic morphology, and autism-like behaviors (Bhattacharya et al., 2012). Given that aberrant mRNA translation is a core pathophysiology of FXS, and that restoring translational homeostasis corrects pathological phenotypes, it is evident that one way to understand the molecular basis of FXS is to (i) identify mistranslated mRNAs, (ii) ascertain the mechanism(s) by which the translation of these mRNAs is altered in FXS, and (iii) establish how these translation mechanisms are corrected in models of FXS rescue.

Several studies have attempted to identify aberrantly translated mRNAs in FXS mice (Ceolin et al., 2017; Das Sharma et al., 2019; Thomson et al., 2017). Intriguingly, all three analyses noted that a large fraction of differentially translated genes showed reduced ribosome association in FXS. Although several explanations have been offered for this puzzling finding, a recent study observing similar reductions in ribosome association with depletion of FMRP in *Drosophila* oocytes led its authors to challenge the dogma that FMRP loss causes increased *de novo* protein synthesis (Greenblatt and Spradling, 2018). There is also limited overlap among the genes identified as aberrantly translated in the studies, necessitating further investigations to clarify the translational targets if FMRP. Moreover, very little is known about the molecular mechanisms by which translational homeostasis is restored through genetic reduction of S6K1 in FXS mice.

In this study, we sought to identify the mRNAs that are mistranslated in FXS mouse brains, ascertain the molecular basis of the alteration in translation, and investigate the mechanism by which genetic deletion of S6K1 normalizes the aberrant translation to levels observed in wild-type (WT) littermates. We carried out ribosome profiling and RNA-Seq on cortical lysates from ~P30 WT, FXS, S6K1 knockout, and double knockout (DKO) mice. Consistent with previous studies, we observed that the majority of genes with differential ribosome association in FXS displayed reduced ribosome footprint abundance. Using molecular assays, we discovered evidence that this reduction reflects a global increase in the rate of ribosome translocation in FXS neurons, is captured by decreased number of translating ribosomes at steady-state, and is normalized by pharmacological inhibition of S6K1. We also observed that alterations in ribosome footprint abundance and mRNA expression show opposing associations with coding sequence (CDS) length in FXS mice, and summate as a positive-to-negative linear gradation in log-fold-changes in translation efficiency (RF/mRNA) with CDS length. Remarkably, this gradation is prevented by the genetic removal of S6K1 in FXS mice. Therefore, we have uncovered the identities of mistranslated mRNAs in FXS, the mechanistic basis of aberrant translation, and a molecular mechanism for correction of *de novo* protein synthesis with genetic reduction of S6K1 in FXS mice.

## Results

To investigate the impact of FMRP loss on the translation of specific mRNAs, we first examined polysome profiles prepared from cortical lysates from P30 FXS mice. We observed little difference in the polysome profiles of FXS mice compared to wild-type littermates (Fig. S1A). We proceeded to perform ribosome profiling to infer mRNA translation profiles at a genome-wide level (Fig. S1B) (Ingolia et al., 2009). The majority (n=153) of the 204 genes with significantly altered ribosome footprint (RF) abundance in FXS cortices showed reduced footprint counts (Fig. 1A), and were enriched for genes linked to glial cell development (Fig. 1F). In stark contrast, of the 351 genes with significantly changed mRNA expression, the majority (n=265) showed elevated expression in FXS brains (Fig. 1A, n=4 FXS, 4 WT), and were enriched for genes implicated in synaptic transmission and development of the nervous system (Fig. 1F).

**Fig. 1.**
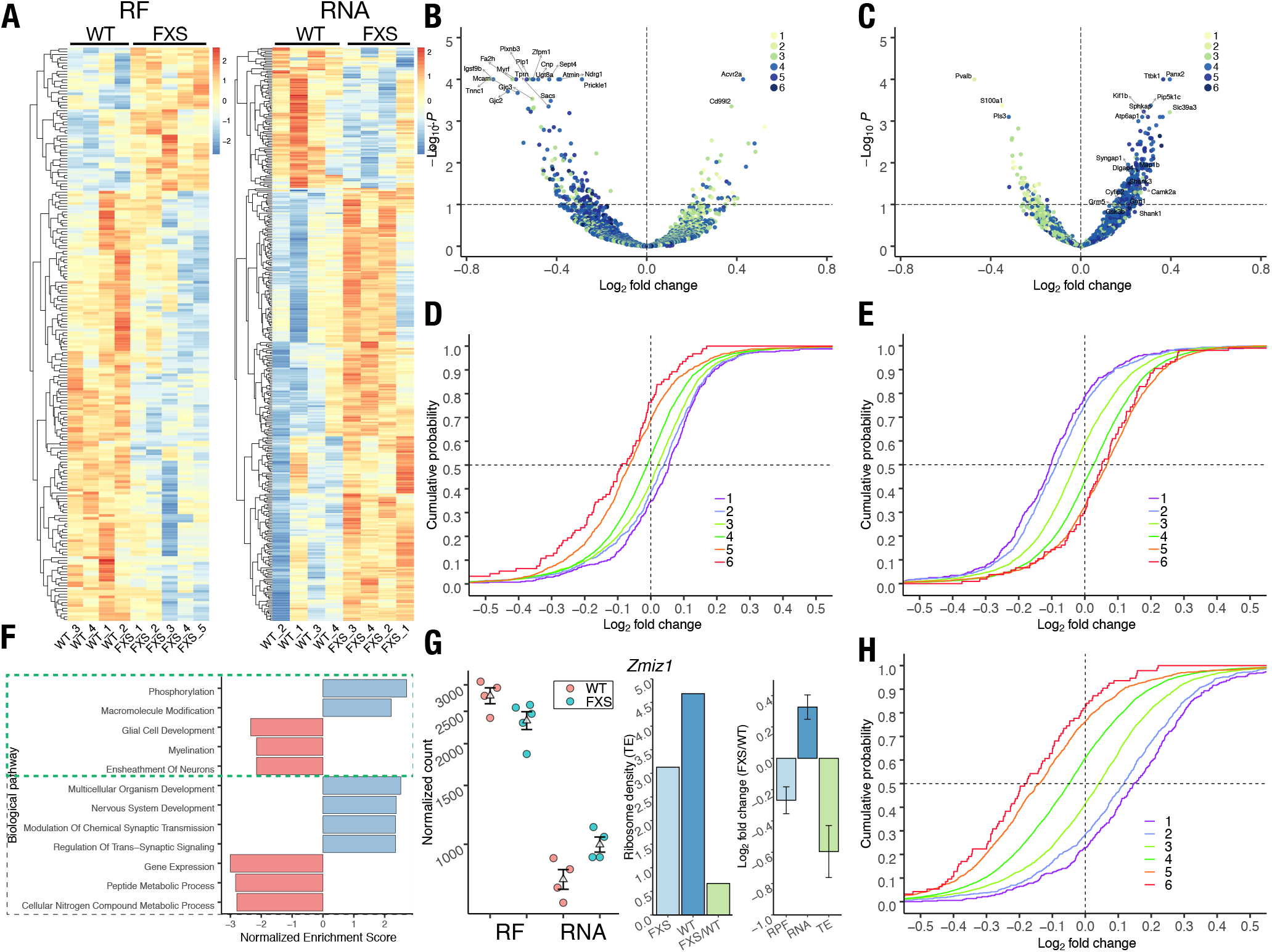
Increased CDS length is associated with decreased ribosome-footprint abundance and increased mRNA expression in FXS mouse cortices. **(A)** Heat maps depicting RF abundance (n=204) and mRNA expression (n=351) of significantly different genes (FDR adjusted p-value< 0.1) between wild-type (WT) and FXS mouse cortices in ribosome profiling (left) and RNA-seq (right) analyses. Each row plots centered and scaled (“row-normalized”) transcripts per million (TPM) values for significantly different genes. **(B)** Significance (FDR adjusted p-value) vs. log-fold change (LFC) in RF abundance between FXS and WT mice. The 20 most significant genes are labeled. **(C)** Volcano plot of mRNA expression between FXS and WT mice. The top 10 significantly different genes, and several genes previously shown to be implicated in FXS are labeled. **(D)** Cumulative distribution of LFCs in RF abundance (FXS/WT) as a function of CDS length. **(E)** Cumulative distribution of LFCs in mRNA expression (FXS/WT) as a function of CDS length. **(F)** Top biological processes enriched in FXS and WT mouse cortical tissue in ribosome profiling (upper green-dashed box) or RNA-seq assays. **(G)** The translation profile of the Zmiz1 gene. The RF abundance LFC (FXS/WT) and the negative of the mRNA expression LFC (FXS/WT) are added to calculate the LFC in translation efficiency. **(H)** Cumulative distribution of LFCs in translation efficiencies (FXS/WT) as a function of CDS length.

The genes with aberrant RF abundance in FXS brains overlapped significantly (p=2.0X10^-3), albeit weakly (n=26), with previously identified binding targets of FMRP (Darnell et al., 2011). In addition, all overlapping genes exhibited reductions in RF abundance in FXS. In contrast, the genes with altered mRNA expression showed a robust overlap with FMRP targets (n=131, p=4.5X10^-66), with almost all (n=129) exhibiting increased mRNA expression in FXS brains.

Previous work has suggested that the translation of mRNAs with long coding sequences (CDSs) is particularly vulnerable to FMRP loss (Greenblatt and Spradling, 2018). To evaluate this phenomenon, we ordered mRNAs ascendingly by their CDS lengths, divided them into six color-coded bins, and evaluated their log-fold changes against their FDR-adjusted p-values (Fig. 1B). Among mRNAs with significantly decreased RF abundance in FXS cortices, we observed enrichment for those with long CDSs. Conversely, mRNAs with short CDSs exhibited a significant upregulation of RFs. Analyzing the fold-changes of all mRNAs revealed that as the CDS length of an mRNA increases, it is more likely to exhibit reduced RF abundance in FXS brains, with the mRNAs harboring the shortest CDSs showing upregulation of footprints (Fig. 1D). Meanwhile, identical analysis of mRNA expression showed opposite associations. mRNAs with long CDSs were upregulated in FXS brains, whereas mRNAs with short CDSs were downregulated (Fig. 1C). Analyzing transcriptome-wide fold-changes revealed that the probability of elevation in mRNA expression was a positive function of CDS length: as the CDS length of mRNA is extended, it was more likely to increase in expression in FXS brains (Fig. 1E). Furthermore, the opposing effects in RF abundance and mRNA expression summate additively to elevate the log-fold changes (LFCs) in translation efficiency (TE) in FXS brains (Fig. 1G, S1C). As expected, evaluating the cumulative probabilities of LFCs in translation efficiency against CDS lengths transcriptome-wide revealed a profile reminiscent to that of RF abundances, but with elevated changes in each CDS length bin (Fig. 1H). We also observed strikingly similar results when we implemented our analysis on a previously published ribosome profiling and RNA-seq dataset from P24 FXS model mouse brains (Fig. S2) (Das Sharma et al., 2019).

A straightforward interpretation of the reduction in RF counts in the majority of altered mRNAs would be that FMRP loss reduces mRNA translation (Greenblatt and Spradling, 2018). However, given evidence suggesting that FMRP directly binds to the ribosome (Chen et al., 2014) and stalls its translocation on specific mRNAs (Darnell et al., 2011), we decided to further investigate the molecular basis of this observation. First, we examined *de novo* protein synthesis in embryonic cortical neurons from FXS mice. To label newly synthesized proteins, we used puromycin, an aminoacyl-tRNA analog that covalently binds nascent peptide chains and causes their premature release. Quantifying these truncated peptides measures *de novo* protein synthesis (Schmidt et al., 2009), which we observed to be elevated in FXS primary neurons (Fig. 2A, 2B).

**Fig. 2.**
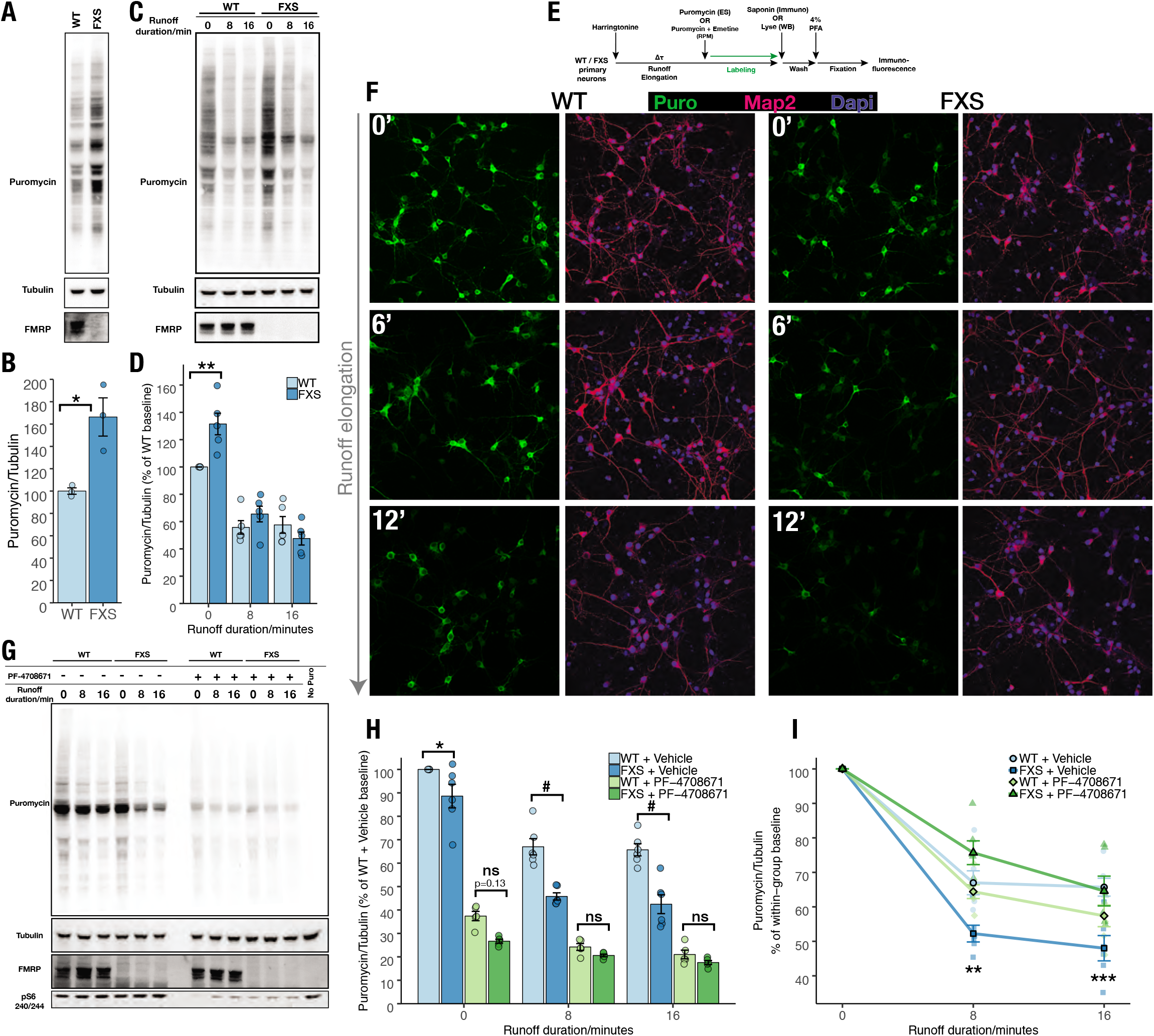
The rate of translation elongation is increased in FXS primary neurons and is sensitive to inhibition of S6K1. **(A)** Representative western blot of WT and FXS cortical neurons incubated with puromycin. **(B)** Quantification of panel A (p=0.05). **(C)** Representative western blot of the elongation SUnSET (ES)assay in WT and FXS neurons. **(D)** Quantification of panel C. (holm-adjusted p=0.007 at baseline). **(E)** Protocol for ES and runoff-ribopuromycylation (runoff-RPM) assays. **(F)** Representative immunofluorescence images of runoff-RPM in WT and FXS mouse cortical cultures. **(G)** Representative western blot of runoff-RPM assay in WT and FXS mouse neurons treated with the S6K1 inhibitor PF-4708671. **(H)** Quantification of panel G. Values are expressed as a percentage of WT + Vehicle baseline. **(I)** The inverse of the rate of runoff elongation. Elongation rate is increased in FXS (t8, WT + Vehicle vs FXS + Vehicle, holm-adjusted p=0.005), and is rescued by pharmacological inhibition of S6K1 (pairwise t-test, t16 WT+Vehicle vs. t16 FXS+PF470861 holm-adjusted p=1).

To examine protein synthesis specifically during the elongation phase of translation, we pulsed FXS neurons with Harringtonine (HHT) for 0, 8, or 16 minutes, then chased them with puromycin (Fig. 2E). HHT inhibits translation initiation rapidly (Gerashchenko et al., 2018). Therefore, after its addition, puromycin labeling specifically measures the elongation phase of protein synthesis. Using this approach, hereafter called “elongation-SUnSET” (ES), we observed elevated elongation-specific protein synthesis at baseline (where HHT and puromycin are added simultaneously) in FXS neurons (Fig. 2C, 2D). Conversely, we observed a decrease in elongation-specific protein synthesis at baseline in FXS mouse embryonic cortical neurons transduced with human homolog of full-length FMRP (Fig. S3A, S3B). These results establish that *de novo* protein synthesis is elevated in FXS neurons even after inhibition of translation initiation, consistent with previous work showing that FMRP loss leads to a global elevation in elongation-specific protein synthesis in cortical lysates (Udagawa et al., 2013). However, one might also expect a similar outcome if FMRP loss led to increased initiation, where mRNAs are already engaged with an increased number of ribosomes before HHT addition.

To examine ribosome engagement specifically, we employed ribopuromycylation (RPM) (David et al., 2013). This assay uses puromycin to tag nascent proteins in the presence of emetine, a translation elongation inhibitor that freezes ribosomes in place on mRNAs without affecting its catalytic activity. Because emetine rescues the ribosome from puromycin-induced dissociation, each translating ribosome is tagged with a single molecule of puromycin. We reasoned that pulsing cultured neurons with HHT and chasing with RPM would permit measurement of the number of translating ribosomes remaining after defined periods of runoff elongation (Fig. 2E). Using this approach, hereafter termed “runoff-RPM,” we observed slightly reduced levels of translating ribosomes in FXS neurons at baseline (Fig. 2F). With increasing runoff durations of 6 and 12 minutes, we observed a faster rate of decrease in the number of translating ribosomes in FXS neurons (Fig. 2F). Conversely, with increased runoff elongation, we observed a slower rate of decrease in the number of translating ribosomes in FXS neurons transduced with human FMRP (Fig. S3C). These results strongly suggest that FMRP impedes ribosome translocation, and that its loss causes a global increase in the rate of translation elongation in neurons.

To quantify elongation kinetics and to better understand how S6K1 regulates translation in FXS, we carried out runoff-RPM followed by western blotting in wild-type and FXS cortical neurons treated with the S6K1 inhibitor PF-4708671. Consistent with the results from immunofluorescence studies, we observed a significant reduction in the number of ribosomes in FXS neurons treated with vehicle at baseline compared to wild-type neurons (Fig. 2G, 2H). This effect was amplified with runoff elongation: fewer translating ribosomes remained after 8 and 16 minutes of runoff elongation in FXS neurons. In contrast, the expression of human FMRP in FXS neurons elevated the number of ribosomes remaining after runoff elongation (Fig. S3E). Inhibition of S6K1 with PF-4708671 dramatically reduced the number of translating ribosomes at baseline in both wild-type and FXS neurons. However, with runoff elongation, the reductions observed in FXS neurons treated with PF-4708671 were not as substantial as those observed in vehicle-treated FXS neurons. This prompted us to determine the rate of loss in the number of translating ribosomes, which we computed by comparing the number of translating ribosomes remaining after runoff elongation in each condition against its own baseline (Fig. 2I, S3F). This metric, which quantifies the inverse of the rate of translation elongation, showed that pharmacological inhibition of S6K1 rescues the excessive rate of translation elongation in FXS neurons. Combined, these results establish that an increase in the rate of ribosome translocation largely mediates elevated *de novo* protein synthesis in FXS neurons. This increase is captured by the reduced number of translating ribosomes with runoff elongation in FXS neurons, which in turn explains why the majority of altered genes in FXS brains have decreased RF counts.

Given our observation that inhibition of S6K1 rescues translation elongation in FXS neurons, we asked whether the previously observed genetic rescue with S6K1 deletion also was mediated by reducing the rate of elongation on specific mRNAs *in vivo*. We examined the RF abundance (n=3) and mRNA expression (n=4) of double knockout (DKO) mice that harbor genetic deletions of both *Fmr1* (FMRP) and *Rps6kb1* (S6K1; Fig. 3A). To our surprise, the DKO mice exhibited significant departures from wild-type mice in both RF abundance (236↑, 355↓; Fig. 3B) and mRNA expression (945↑, 666↓; Fig. 3C) for a large number of genes. However, we did not observe any clear CDS-length dependency among mRNAs with significantly altered RF abundance or mRNA expression in DKO brains. We confirmed this transcriptome-wide by analyzing cumulative probabilities against LFCs of RF abundance (Fig. 3D) and mRNA expression (Fig. 3E) in DKO cortices. To compare DKO mice directly to their FXS littermates, we ordered mRNAs into 50 bins by their CDS lengths and examined their fold changes compared to wild-type mice. We observed that genetic reduction of S6K1 in FXS mice specifically normalized their aberrant deviations in mRNA expression (Fig. 3F) and translation efficiency (Fig. 3G, S4A), particularly for genes with especially long or short CDSs. Furthermore, examining the RF abundance and mRNA expression in FXS and DKO brains of the genes in the final four bins revealed that genetic reduction of S6K1 normalizes gene expression by moving the RF abundance (Fig. 3H), and especially mRNA expression (Fig. 3I), of the entire bulk of genes towards the wild-type, rather than by rescuing specific genes. Identical evaluation of genes in the first four bins also showed a strikingly concordant effect (Fig. S4B, S4C).

**Fig. 3.**
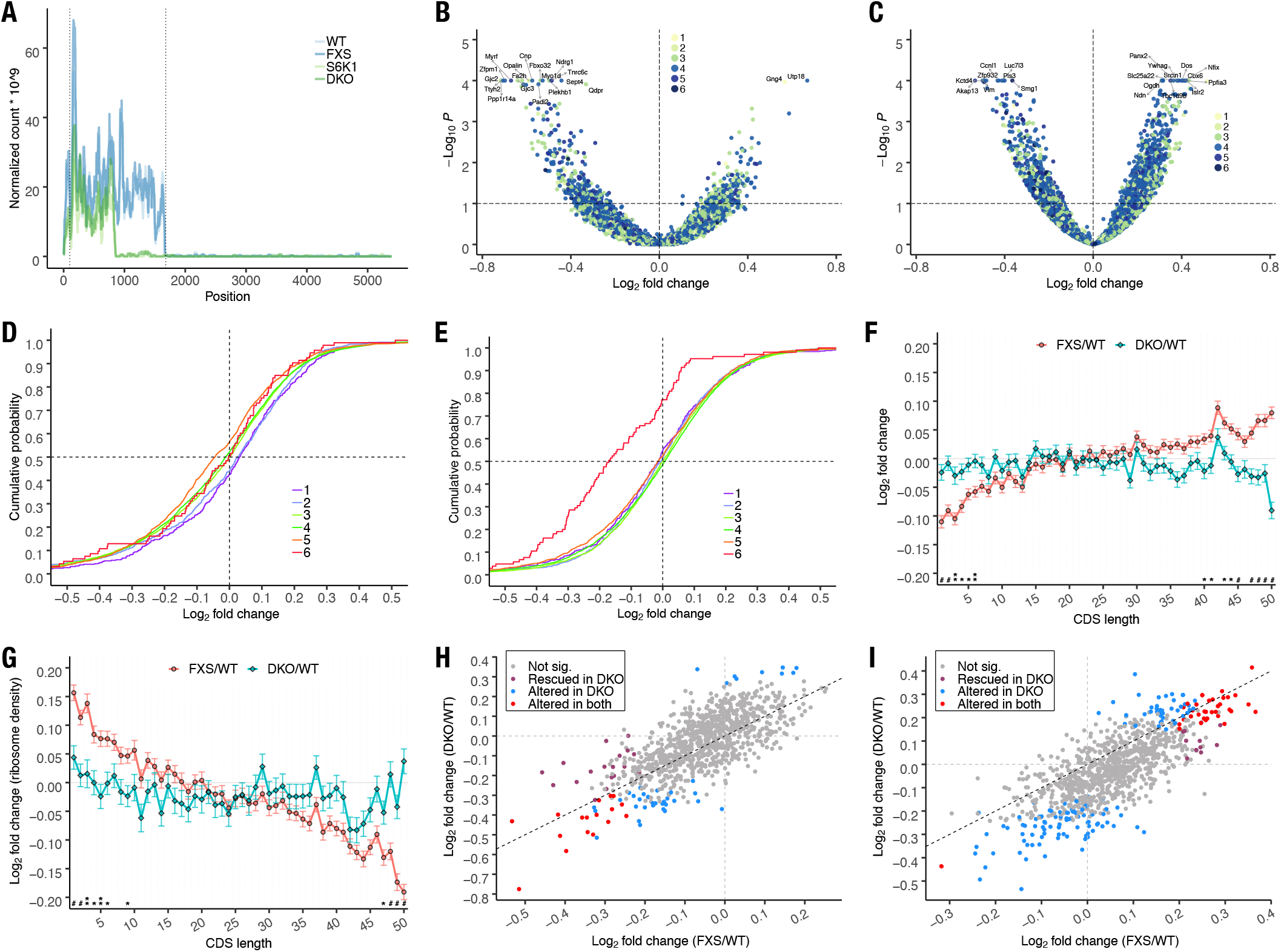
CDS length-dependency of translation efficiency and mRNA expression in FXS mouse cortices is corrected by genetic reduction of S6K1. **(A)** Ribosome footprint profile of Rps6kb1 mRNA. Profile accurately captures the stop codon introduced to make the S6K1 deletion. Dotted lines indicate untranslated region (UTR) boundaries. **(B)** Volcano plot of RF abundance (n=3) and **(C)** mRNA expression (n=4) (DKO/WT). The top 20 genes are labeled. **(D)** Cumulative distribution of LFCs in RF abundance and **(E)** mRNA expression in DKO brains, as a function of CDS length. **(F)** Comparison of LFCs by CDS length in FXS and DKO brains in mRNA expression and **(G)** translation efficiency datasets. **(H)** Comparison of RF abundance and **(I)** mRNA expression in FXS and DKO brains of genes in the longest 4 bins, compared to the WT. Majority of points are above the 45-degree line in RF abundance and below the line in RNA expression, suggesting a generalized reduction in RF abundance and mRNA expression towards WT levels.

## Discussion

The translation efficiency of an mRNA historically has been deduced by the number of polysomes bound to it (Arava et al., 2003). Remarkably, an inverse correlation between CDS length and translation efficiency - the longer the CDS length of a transcript, the lower its ribosome density - has been observed in multiple model systems ranging from the protozoan *P. falciparum* to human cells (Fernandes et al., 2017). This relationship therefore suggests a CDS length-dependent effect on translational control which might be shared across phyla.

Because each ribosome footprint reports on a single ribosome bound to an mRNA, the total number of RF reads aligning to a transcript directly reflects the overall number of polysomes bound to that mRNA species. Using ribosome profiling and RNA-Seq, we uncovered a positive-to-negative gradation of alterations in translation efficiency in the cortices of FXS model mice. This gradation suggests that the mechanisms regulating CDS length-dependent control of translation go awry with the loss of FMRP.

How does CDS length control translation? Using an elegant mathematical model, Fernandes et al. (2017) propose that mRNAs with short CDSs exhibit an increased rate of translation initiation. They assume that the local concentration of ribosomal subunits around the 5’ UTR of an mRNA determines its initiation rate. The shorter the CDS, the closer the 5’ end of an mRNA is to its 3’ end. After translation terminates, the concentration of ribosomal subunits at the 5’ UTR is therefore higher for short CDS mRNAs, which increases the initiation rate on those mRNAs. Ingolia et al. (2009), meanwhile, propose that mRNAs with long CDSs exhibit an increase in the rate of translation elongation. Consistent with previous polysome profiling studies, these researchers observed that shorter genes tended to have higher ribosome densities. Examination of the average ribosome densities of thousands of mRNAs revealed a high ribosome density towards the 5’ end of their CDSs, which tapered off and persisted at a uniform level towards the 3’ end. This trend suggested that the reduction in ribosome density with CDS length may be mediated by an increase in the rate of elongation on distal parts of the transcript on long CDS mRNAs (Ingolia et al., 2009). Therefore, in the absence of any exogenous changes, mRNAs with short CDSs are likely to exhibit increased rates of initiation, while those with long CDSs are likely to exhibit increased rates of elongation.

If these two proposals are true, what would be the consequences if FMRP were genetically removed from this system? Using molecular assays, we have provided substantial evidence that the rate of translation elongation is globally elevated in FXS. FMRP loss therefore likely causes elevated rates of elongation on mRNAs with short or long CDSs. On mRNAs with especially short CDSs, the increase in the rate of elongation would in turn increase the rate of termination, and subsequently elevate the local concentration of ribosome subunits around their 5’ UTRs. This increase would cause further enhancement in the rate of translation initiation on those mRNAs. On mRNAs with long CDSs, the elevated rate of elongation would cause additional reductions in ribosome density. Importantly, both of these effects would be determined directly by CDS length. This model would therefore predict that loss of FMRP causes an elevation in the TEs of mRNAs with short CDSs, a reduction in the TEs of mRNAs with long CDSs, and a CDS length-dependent gradation between the two extremes. Remarkably, the predictions of this model exactly match the alterations we observed in the cortices of FXS mice. The negative-to-positive linear gradation of alterations in RF abundance and TE may therefore be an emergent property of an increased rate of ribosome translocation in FXS.

Because FMRP is thought to bind only to ~1000 mRNAs (Darnell et al., 2011), it is also possible that the global increase in elongation rate is caused by a generalized increase in ribosome translocation due to hyperactivation of mammalian target of rapamycin complex 1 (mTORC1) to S6K1 signaling, which has been previously noted in FXS brains (Sharma et al., 2010). This is consistent with the limited, albeit significant, overlap between genes with altered RF abundance and FMRP binding targets, and the reduction in the rate of translation elongation with inhibition of S6K1 in FXS neurons. Meanwhile, increased translation initiation due to elevated mTORC1 signaling to its other major downstream targets (4E-BP and/or CYFIP1), and resulting increases in eIF4F formation (Napoli et al., 2008), explain the increase in footprint expression in the shortest genes. These genes are highly enriched in 5’ terminal oligopyrimidine mRNAs which code for ribosomal proteins, and are established translational targets of mTORC1 activation (Fig. S5) (Thoreen et al., 2012). Indeed, the positive-to-negative gradation in RF abundance and translation efficiency by CDS length argues for a model where elevated rates of both initiation and elongation act in concert to elevate mRNA translation indiscriminately in FXS brains. We hypothesize that precisely because initiation is rate-limiting, its increase is best captured in mRNAs with short CDSs. Meanwhile, the higher rate of elongation dominates in longer transcripts because ribosomes must translocate through a greater number of codons, with the net effect of each increased translocation event aggregating over the length of the CDS. Our proposed models are consistent with and provide mechanistic insights into previously proposed models of FMRP function (Das Sharma et al., 2019; Chen et al., 2014; Sharma et al., 2010; Napoli et al., 2008). Alternatively, it is possible that FMRP binding is more promiscuous than appreciated and biased towards longer mRNAs, which are enriched in neurons (Zylka et al., 2015). Therefore, the increased rate of elongation is primarily observed in longer transcripts, and the secondary effect of increased rate of initiation is observed in shorter transcripts, which are translational targets of mTORC1 (Fig. S5B).

Elevated rates of ribosome translocation also have been associated with increased mRNA stability (Hanson et al., 2018). Specifically, an increase in the rate of decoding of cognate transfer RNAs at the ribosomal A-site is thought to drive increases in mRNA stability. Because this process is likely elevated in every codon in FXS (Das Sharma et al., 2019), the effect on mRNA stability is naturally summated over the length of a transcript. This model accounts for the increased expression of mRNAs with long CDSs in FXS brains, as well as the negative-to-positive gradation in mRNA expression by CDS length. Finally, based on our *ex vivo* findings with inhibition of S6K1, we propose that genetic reduction of S6K1 reduces translation elongation in FXS mice. This, in turn, decreases mRNA stability, which is particularly summated over long genes, thereby driving the rescue of expression towards the wild-type.

In addition to establishing bona fide targets for pharmacological interventions in FXS, our results ascertain the underlying biological principles of FMRP function in health and disease. Our work also suggests that drugs that reduce translation elongation on specific target mRNAs (Li et al., 2019) may offer a novel therapeutic avenue in FXS.

## Materials and Methods

### Mouse breeding

DKO mice were generated initially by crossing heterozygous female FXS mice carrying the *Fmr1* mutation with heterozygous male mice carrying the S6K1 mutation. Subsequently, all animals used for experimentation were derived from the crossing of female X^Fmr1^X^+^;S6K1/+ with either X^+^Y;S6K1/+ or X^Fmr1^Y;S6K1/+ males (Bhattacharya et al., 2012). All experimental animals were littermates. All procedures involving animals were performed in accordance with protocols approved by the New York University Animal Welfare Committee and followed the National Institutes of Health (NIH) Guide for the Care and Use of Laboratory Animals. All mice were housed in the New York University animal facility and were compliant with the NIH Guide for the Care and Use of Laboratory Animals. Mice were housed with their littermates in groups of two to three animals per cage and kept on a 12-hour regular light/dark cycle, with food and water provided ad libitum.

### Polysome profiling

Cortices from P29-34 mice were rapidly dissected post cervical-dislocation and washed with ice-cold 10mM Hepes-HCl in HBSS. The tissue then was dounce-homogenized (40 strokes; Corning 7724T3) in 1ml of lysis buffer consisting of 25mM Hepes-HCl pH 7.3, 150mM KCl, 5mM MgCl_2_, 0.5mM dithiothreitol (DTT), 100μgml^−1^ Cycloheximide, 10Uml^−1^ Superase-In, and 1X Halt protease and phosphatase inhibitor. The lysate was centrifuged at 2000g (4°C) for 10 minutes and NP-40 (Sigma I8896) was added to a final concentration of 1% to the supernatant, which then was centrifuged at 10,000g (4°C) for 5 minutes. The final supernatant was gently layered onto 10-50% sucrose density gradients prepared in 25mM Hepes-HCl pH 7.3, 150mM KCl, 5mM MgCl_2_, 0.5mM DTT and 10Uml^−1^ SuperaseIn. The gradients were ultracentrifuged at 35000 rpm on a SW41-Ti rotor for 2.5 hours (4°C). To obtain polysome profiles, gradients then were fractionated with continuous monitoring of ultraviolet absorbance at 254nm in a Biocomp Piston Gradient Fractionator.

### Isolation of ribosome footprints and total RNA

Cortices from P29-34 WT (n=4), FXS (n=5), S6K1 (n=3) and DKO (n=3) animals were rapidly dissected, washed with ice-cold 10mM Hepes-HCl pH 7.3 in HBSS, then lysed by dounce homogenization (40 strokes) in 25mM Hepes-HCl pH 7.3, 150mM KCl, 5mM MgCl_2_, 0.5mM DTT, 100μgml^−1^ Cycloheximide, and protease/phosphatase inhibitors on ice. Lysates were centrifuged at 2000g for 10 minutes (4°C) to remove cellular debris and nuclei. Ten A260 units of the supernatant was incubated with 1 ug RNase A and 600 units RNase T1 with gentle rotation for 30 minutes at room temperature. RNase digestion was stopped by placing the tubes on ice subsequent to the addition 600 U Superase-In to each reaction. The RNase digested lysates then were supplemented with NP-40 to a final concentration of 1 %. After incubating on ice for 5 minutes, the lysates were centrifuged at 10,000g for 5 minutes, and the supernatants layered on a 10-50% sucrose density gradient prepared in 25mM Hepes-HCl pH 7.3, 150mM KCl, 5mM MgCl_2_, 0.5mM DTT and 10Uml^−1^ Superase-In. The gradients were ultracentrifuged for 2.5 hours (4°C) in a SW41Ti rotor (Beckman Coulter) at 35000 rpm, after which they were fractionated on a Biocomp Piston Gradient Fractionator with continuous UV absorbance monitoring (254nm) on the EM1 UV Monitor (Bio-Rad). Monosome fractions were isolated and RNA extracted with Trizol-LS. For total RNA isolation (n=4 each of WT, FXS, S6K1 and DKO), cortices were lysed as described. Lysates were supplemented with 600U Superase-In and 1% NP-40, the RNase digestion step skipped, and 10,000g supernatant isolated. The cortex from a single mouse was used to prepare one ribosome footprint or RNA-seq high-throughput sequencing library.

### High-throughput sequencing library preparation

Ribosome footprints were converted to sequencing libraries as described previously (Cenik et al., 2015). Briefly, ribosomal RNA was depleted from monosome fractions with Ribo-Zero Gold, after which 26-34 nt long monosomal RNA was fractionated out using a 15% Urea-Page gel. The excised RNA was 3’ de-phosphorylated, ligated to a pre-adenylated adapter, reverse-transcribed, circularized and PCR amplified to generate strand-specific sequencing libraries. For RNA-seq, the TruSeq stranded mRNA kit (Illumina) was employed to select polyA-tailed RNAs and convert them to strand-specific sequencing libraries.

### High-throughput sequencing data processing

Ribosome profiling reads were demultiplexed using in-house scripts. Sequencing adapters were removed using CutAdapt. Reads aligning to ribosomal and transfer RNA were excised using Bowtie2. The remaining reads were aligned to the mouse genome (mm10) using TopHat2, with segment length set at 12. PCR duplicates were removed from alignments using unique molecular identifier (UMI) information present in the reads. All reads with non-unique alignments were also discarded. Ribosome footprint abundance was calculated by directly aligning the remaining reads specifically to the coding sequence (CDS) region of *Mus musculus* protein-coding genes using RSEM. To calculate mRNA expression deriving from CDS-derived reads, RNAseq reads were processed identically, except removal of duplicate reads was not carried out because of the lack of UMI information in RNAseq libraries.

To construct the ribosome footprint profile of an individual mRNA, all 5’ alignments at each position along the mRNA was obtained with Plastid (Dunn and Weissman, 2016). The total number of alignments at each position derived from a given library was normalized by the size factor computed by *DESeq2* for that library. Rolling means of the average normalized counts of all samples of the same genotype were plotted to reveal the average ribosome footprint profile of the mRNA.

### Statistical analysis of RF expression and mRNA expression

*DESeq2* (Love et al., 2014) was utilized to model the RF and RNA-seq data. Gene level expression/abundance estimated by RSEM for RF and RNA-seq were collated into respective count matrices and used as input for *DESeq2*. Following the authors’ recommendations, we modeled data from all four genotypes together. Genes with less than 1 count per million in any of libraries were excluded from the analysis. The threshold for differential expression was set at a False Discovery Rate (FDR) of 0.1.

### Gene set enrichment analysis

We used the R package *fgsea* to carry out gene set enrichment analysis (Sergushichev, 2016). Genes were pre-ranked by the Wald test statistic computed when testing for differential expression in *DESeq2*. Only significantly different genes in the RF or RNAseq assays at FDR<0.1 were used in this analysis.

### Gene set overlap analysis

We used the hypergeometric test to calculate the probability that the genes differentially expressed in FXS cortices in either RF or mRNA expressions overlapped by chance with the genes thought to be binding targets of FMRP. Specifically, we used the *phyper* function in R, with the total number of genes set by the number of genes that were included in the *DESeq2* analysis (12083 for RF and 12876 for RNA-seq).

### Division of mRNAs into CDS length bins

Annotations for all *Mus musculus* protein-coding genes in RefSeq were obtained from UCSC table browser. Using these annotations, the CDS region of the mRNAs was extracted using in-house scripts. This truncated transcriptome was used to prepare references for RSEM. When calculating expression, RSEM summates the length of all mRNA isoforms of the gene into an ‘effective length’ for that gene. The effective lengths were log-transformed, and its histogram was divided at regular intervals to categorize genes into 6 CDS length bins. To divide mRNAs into 50 bins evenly, the total number of robustly expresed mRNAs (counts per million > 1 for all samples) were divided by 50 and the number of genes indicated by the remainder was removed at random. For example, 33 genes were removed at random from 12033 genes to ensure that each length bin contained 240 mRNAs.

### Primary neuron culture

Primary neuron cultures were obtained from E16.5 WT and FXS mice. Cultures were prepared as previously described (Myrick et al., 2014). Briefly, dams were sacrificed by CO_2_ inhalation in accordance with the Institutional Animal Care and Use Committee guidelines. Embryos were rapidly isolated into ice-cold HBSS supplemented with Glucose, and their cortices and hippocampi dissected out. The embryonic tissue was trypsinized, triturated, and plated onto either cell culture dishes or coverslips coated with Poly-D-Lysine and Laminin. Neurons were cultured in Neurobasal media supplemented with B27, Glutamax and Penicillin/Streptomycin in a 5%CO_2_ incubator at 37°C. Fresh media was supplemented every three days. All experiments were carried out on DIV 7-8 neurons. To keep the environment constant, WT and FXS cultures were plated on different wells of the same multi-well plate, and all experiments were carried out simultaneously for WT and FXS neurons.

### Lentiviral transduction

Lentivirus constitutively expressing full length N-terminal Flag tagged human *FMR1* and a control lentivirus constitutively expressing EGFP were produced at Emory Viral Vector Core. Both constructs are cloned into the pFUGW plasmid (Myrick et al., 2014). Infection was carried out by incubating DIV2 neurons with lentivirus diluted in conditioned media for 24 hours, after which the viral media was replaced with previously-saved conditioned media.

### Measurement of *de-novo* protein synthesis (SUnSET)

DIV 7-8 WT and FXS primary neurons were treated with 0.5 μgml^−1^ Puromycin in conditioned media for 15 minutes. Cells then were washed once with ice-cold PBS and lysed in RIPA buffer supplemented with 1X Halt protease and phosphatase inhibitor. Lysates were flash-frozen, processed as described, and analyzed on Western blots.

### Elongation SUnSET (ES)

DIV 7-8 WT and FXS primary neurons were treated with 5μgml^−1^ Harringtonine to induce runoff elongation. After 0, 8, or 16 minutes of treatment, the runoff media was replaced with a mix of 5 μgml^−1^ Harringtonine and 50μgml^−1^ Puromycin for 15 minutes to continue inhibiting translation initiation while simultaneously measuring the functional output of only the elongation phase of protein synthesis. Cells then were washed with ice-cold PBS and lysed in RIPA buffer supplemented with 1X Halt. All treatments were carried out in 50% conditioned media. Lysates were flash-frozen, processed as described, and analyzed on Western blots.

### Runoff-ribopuromycylation (Runoff-RPM)

DIV 7-8 WT and FXS primary neurons were treated either with Vehicle (0.1% DMSO) or 10μgml^−1^ PF-4708671, diluted from a freshly prepared 10mgml^−1^ stock in DMSO. After 1 hour of drug treatment, runoff elongation was carried out for 0, 8, or 16 minutes by replacing the drug-treatment media with runoff media, which consisted of 5μgml^−1^ Harringtonine and Vehicle/ 10μgml^−1^ PF-4708671. At the end of each time-point, runoff media was aspirated out and replaced with labeling media made of 5μgml^−1^ Harringtonine, 200μM Emetine, 50μgml^−1^ Puromycin, and Vehicle/10μgml^−1^ PF-4708671. Labeling was carried out for 5 minutes, after which neurons were washed with ice-cold PBS and lysed with RIPA buffer supplemented with 1X Halt. Lysates were flash-frozen, processed as described, and analyzed on Western blots. The initial S6K1 inhibitor treatment was carried out in 100% conditioned media, and all other treatments were carried out in 50% conditioned media.

For imaging, experiments were carried out identically, except after labeling, coverslips with neurons were washed with room-temperature 0.004% Saponin + 5μgml^−1^ Harringtonine + 200μM Emetine in HBSS for 1 minute to pre-permeabilize the cells. The coverslips then were washed rapidly with warm (~37°C) HBSS + 5μgml^−1^ Harringtonine + 200μM Emetine twice to remove unbound puromycin, after which they were fixed with 4% PFA in PBS for 20 minutes at room-temperature.

### Western blotting

Flash-frozen lysates were thawed and briefly sonicated on ice, and centrifuged at 20,000g for 10 minutes (4°C). The supernatant was saved and its protein concentration was measured using the BCA assay. 6X Laemmli-buffer (Boston Bioproducts #BP-111R) was added to the sample and mixed thoroughly, after which the sample was boiled for 6 minutes. 15-20ug of protein from each sample was loaded on 4-12% Bolt Bis-Tris gradient gels (Invitrogen), followed by semi-dry transfer onto nitrocellulose membranes. Membranes were blocked for 90 minutes with 5% milk in Tris-buffered saline supplemented with 0.1% Tween-20 (TBST). Membranes were then probed overnight (4°C) with 1:1000 anti-Puromycin antibody (MABE343, Millipore) in 5% milk TBST with gentle rotation. After 4 vigorous 5 minute washes in TBST, the membranes were incubated with 1:10,000 HRP conjugated anti-mouse IgG antibody for 1 hour in 5% milk TBST at room temperature. Membranes then were washed again 4 times vigorously in TBST, and imaged on the FluorChem-E platform (Protein Simple). ECL and exposures were set to obtain signals in the linear range. Membranes were then stripped, re-blocked with 5% milk TBST, and cut into three pieces with horizontal incisions at ~60 kDA and ~37 kDA. The top portion of the membrane was probed with 1:1000 anti-FMRP antibody (Biolegend 834601), the middle portion with 1:1000 anti-γ-Tubulin (Sigma T5326), and the bottom with 1:1000 anti-phospho S6 Ser 240/244 antibody (Cell Signaling 5364), all in 5% milk TBST. Membranes were imaged for the respective antibodies again as described.

### Microscopy

Coverslip grown neurons fixed after runoff-RPM were washed with PBS twice, permeabilized with 0.2% Triton-X in PBS for 5 minutes, and blocked with 5% BSA in PBS for 30 minutes. Coverslips were incubated with 1:1000 anti-Puromycin conjugated to Alexa Fluor 488 (Millipore MABE343-AF488), and 1:500 Ch. anti-Map2 (Encor Biotech CPCA-MAP2) overnight (4°C) in 1% BSA in PBS. Subsequent to three 5 minute PBS washes, coverslips were probed with 1:200 anti-Chicken IgG conjugated with Alexa fluor 647 in 1% BSA for 1 hour at room-temperature. After another 3 washes in PBS, coverslips were mounted and imaged on a Leica SP8 confocal microscope. All microscopy parameters including laser intensities, gains, and offsets were optimized on WT baseline (0 minutes of runoff), after which every coverslip of either genotype was imaged using the exact same settings.

## Acknowledgments

We thank Thomas Dever (NIH) for a critical reading of our manuscript, Leila Myrick and Stephen T. Warren (Emory) for generously providing the human full-length FMRP expression plasmids, and Botao Liu, Huan Shu, Elisa Donnard, and Joel Richter (UMassMed) for a demonstration of, and technical advice on ribosome profiling library preparation and data analysis.

## Author contributions

E.K. conceived the ribosome profiling experiments. S.A. conceived the translation elongation experiments. S.A. conducted the experiments and analyzed the data, with input from F.L. and E.K. S.A. and E.K. wrote the manuscript.

## Data availability

All data used in the analysis has been deposited under record number GSE143659 at NCBI’s Gene Expression Omnibus (GEO) database.

---

**Fig. S1.**
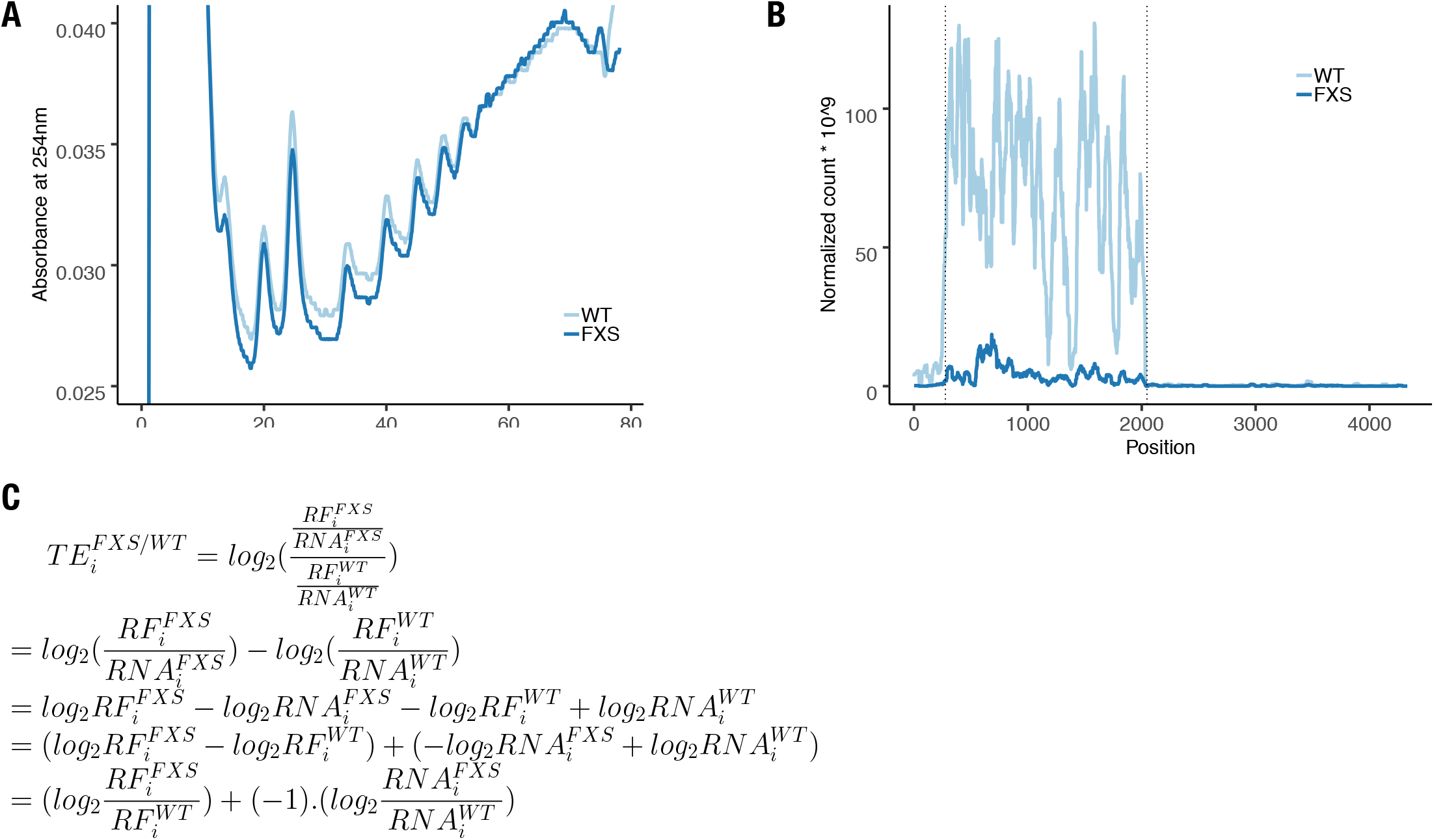
Translation profiling of FXS model mice. **(A)** Polysome profiles of cortical tissue from P30 wild-type (WT) and FXS mouse brains. **(B)** Ribosome profile of *Fmr1* mRNA. Dotted lines indicate untranslated region (UTR) boundaries. **(C)** Simple rearrangement of terms reveals that log-fold-changes in translation efficiency for a gene *i* in FXS can be computed by subtracting its log-fold-change in RNA expression in FXS from the log-fold-change in ribosome footprint in FXS.

**Fig. S2.**
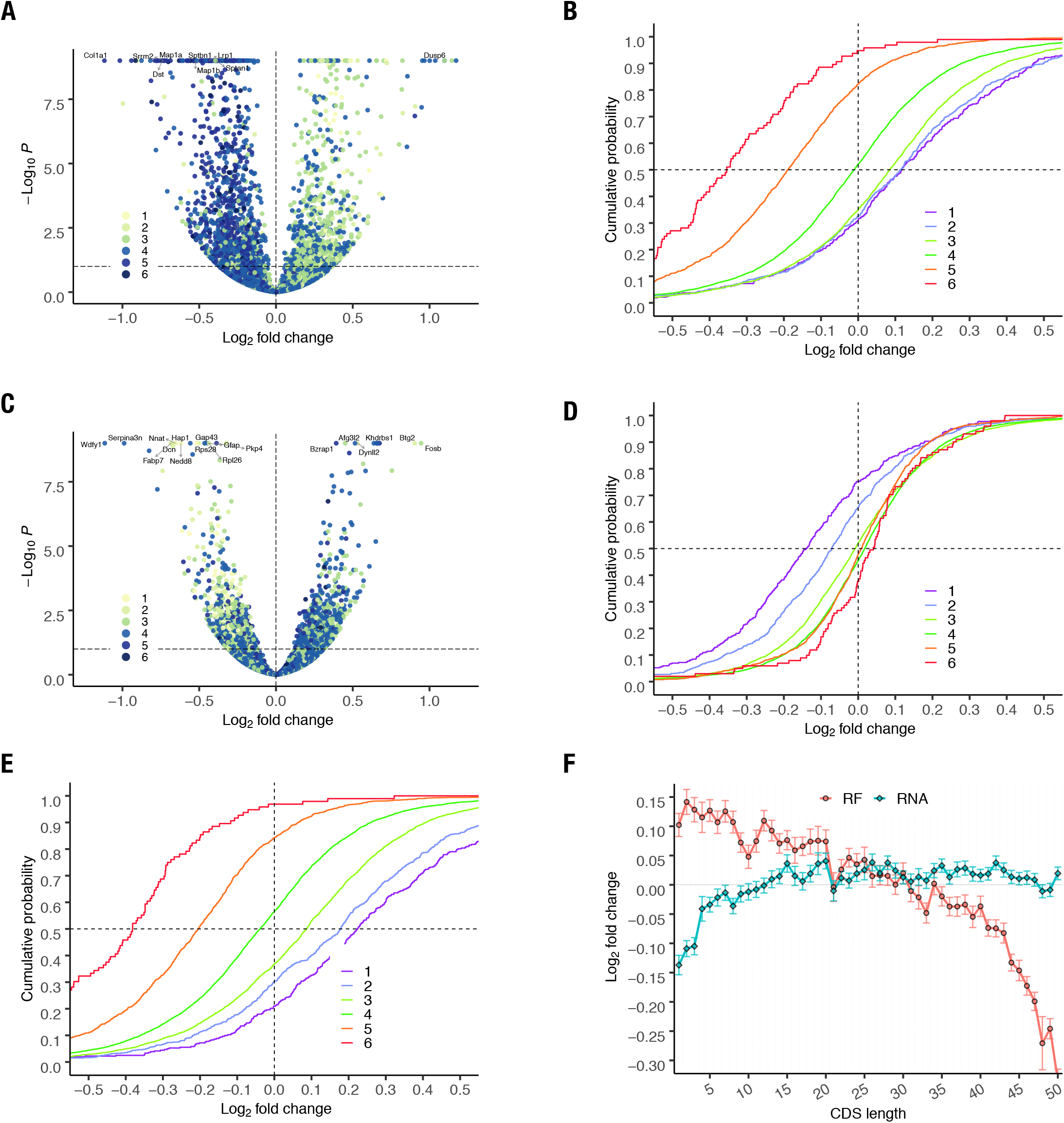
Re-analysis of data from Das Sharma et. al. (2019) also shows that alterations in ribosome footprint abundance and mRNA expression in P24 FXS brains are positive and negative functions of CDS length. **(A)** Volcano plot of RF abundance and **(C)** RNA expression in FXS brains. **(B)** Cumulative distribution of LFCs in RF abundance, **(D)** RNA expression, and **(E)** translation efficiency in FXS brains, as a function of CDS length. **(F)** Log-fold-changes in RNA expression and RF abundance by CDS length. mRNAs were ordered ascendingly by their CDS length and divided into 50 bins.

**Fig. S3.**
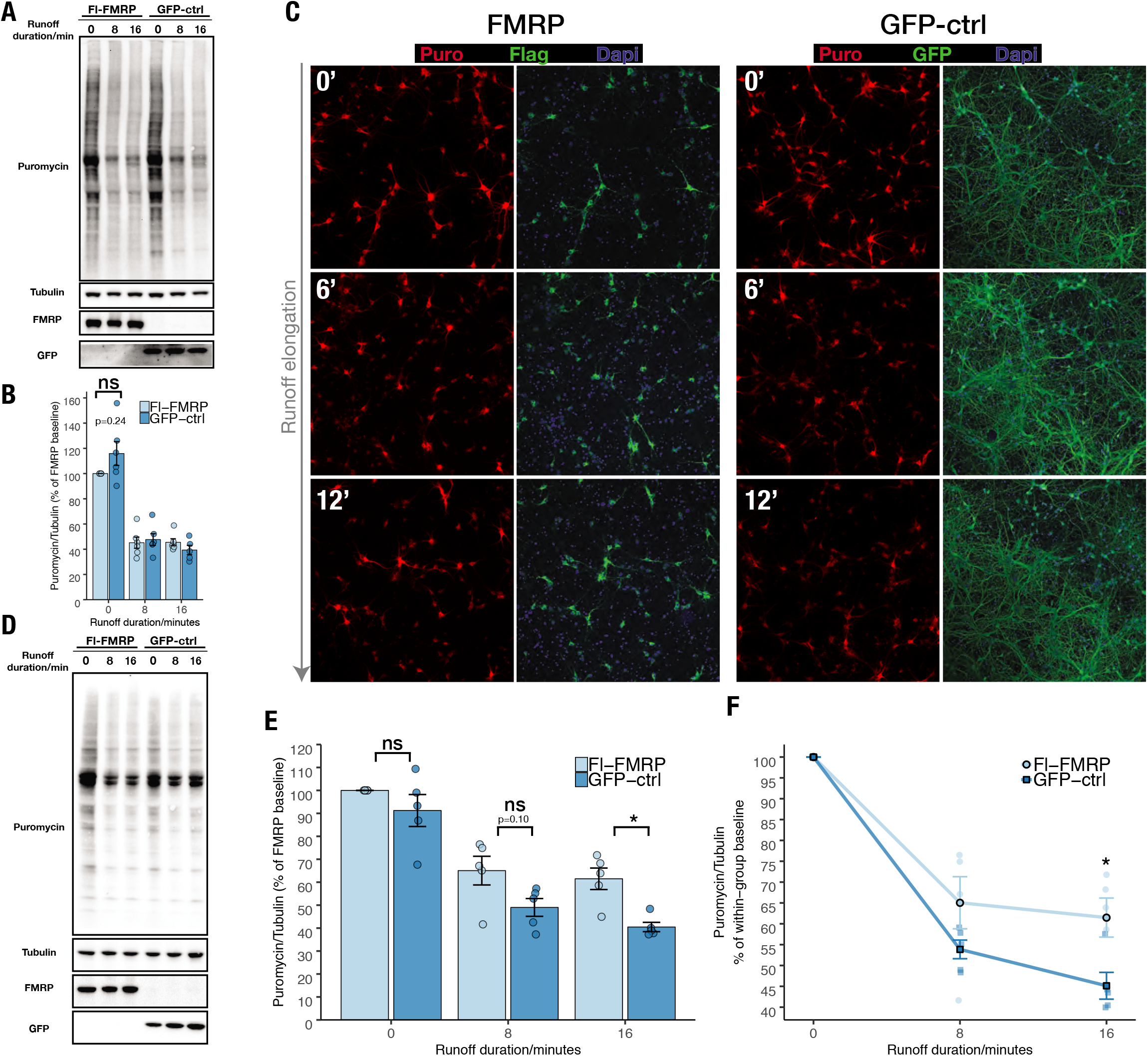
Transduction of human FMRP in FXS primary neurons decreases the rate of translation elongation. **(A)** Representative western blot of elongation SUnSET (ES) carried out in FXS neurons transduced with lentivirus expressing flag-tagged human FMRP isoform 13 (Fl-FMRP) or GFP expressing control (GFP-ctrl) in pFugW backbone. **(B)** Quantification of panel A. Transduction of Fl-FMRP reduces elongation-specific protein synthesis (holm-adjusted p-value = 0.24). **(C)** Runoff-ribopuromycylation in FXS cortical cultures transduced with human FMRP or GFP control. With increased duration of runoff elongation, FXS neurons expressing FMRP exhibit increased ribopuromycylation signal compared to those expressing GFP control. **(D)** Representative western blot of runoff-ribopuromycylation assay carried out in FXS neurons transduced with FMRP or GFP control. **(E)** Quantification of panel D. The number of ribosomes after 16 minutes of runoff elongation is significantly increased in FXS neurons transduced with Fl-FMRP (holm-adjusted p=0.02). **(F)** Results from runoff-ribopuromycylation expressed relative to its own group’s baseline. Expression of Fl-FMRP in FXS neurons significantly reduces the rate of signal loss (t16; p=0.02), therefore decreasing translation elongation.

**Fig. S4.**
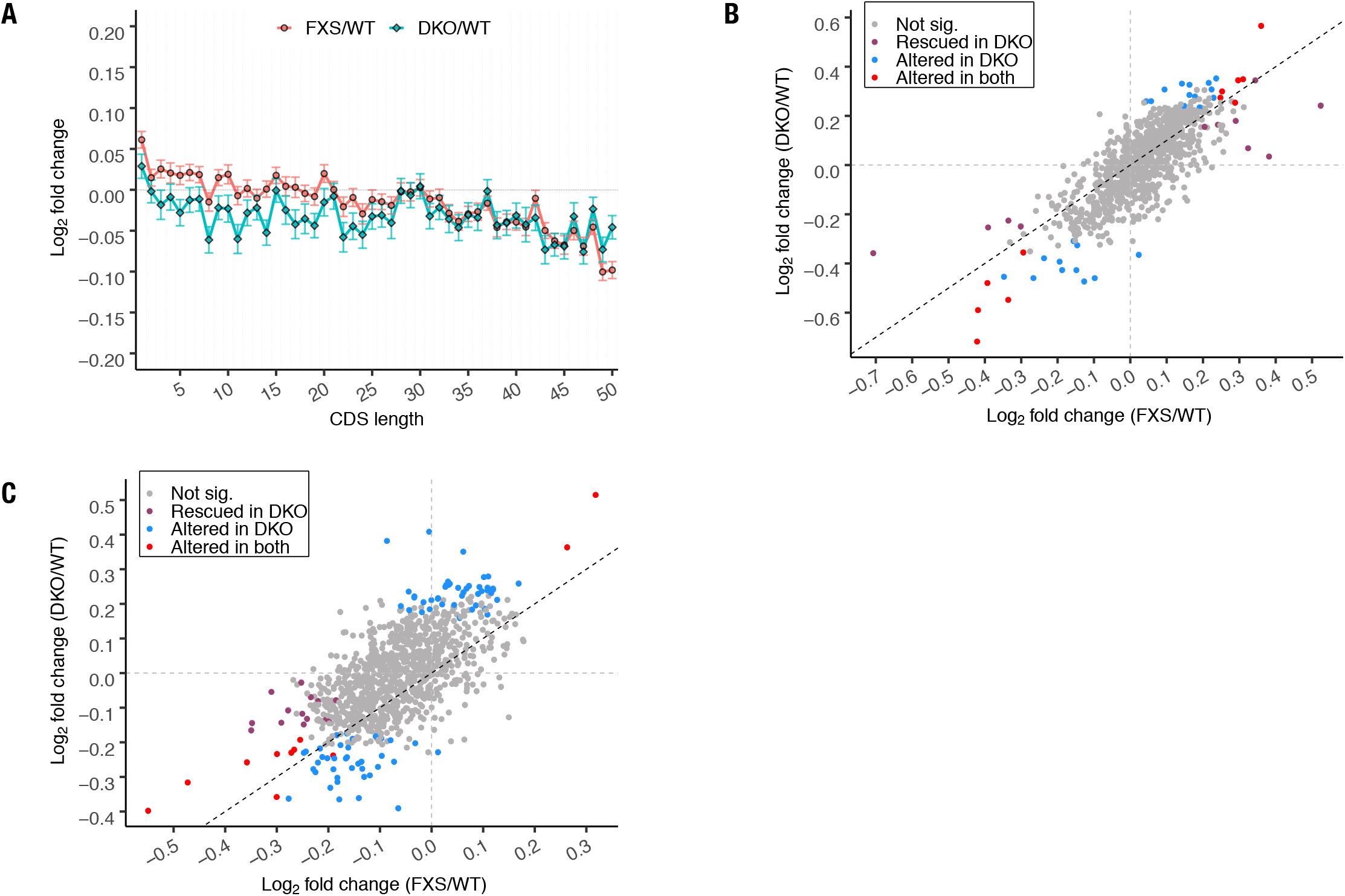
Genetic deletion of S6K1 normalizes overall ribosome footprint abundance and mRNA expression of short genes. **(A)** Comparison of LFCs in RF abundance by CDS length in FXS and DKO brains **(B)** Evaluation of RF abundance and **(C)** mRNA expression in FXS and DKO brains of genes in the shortest 4 bins, compared to the WT.

**Fig. S5.**
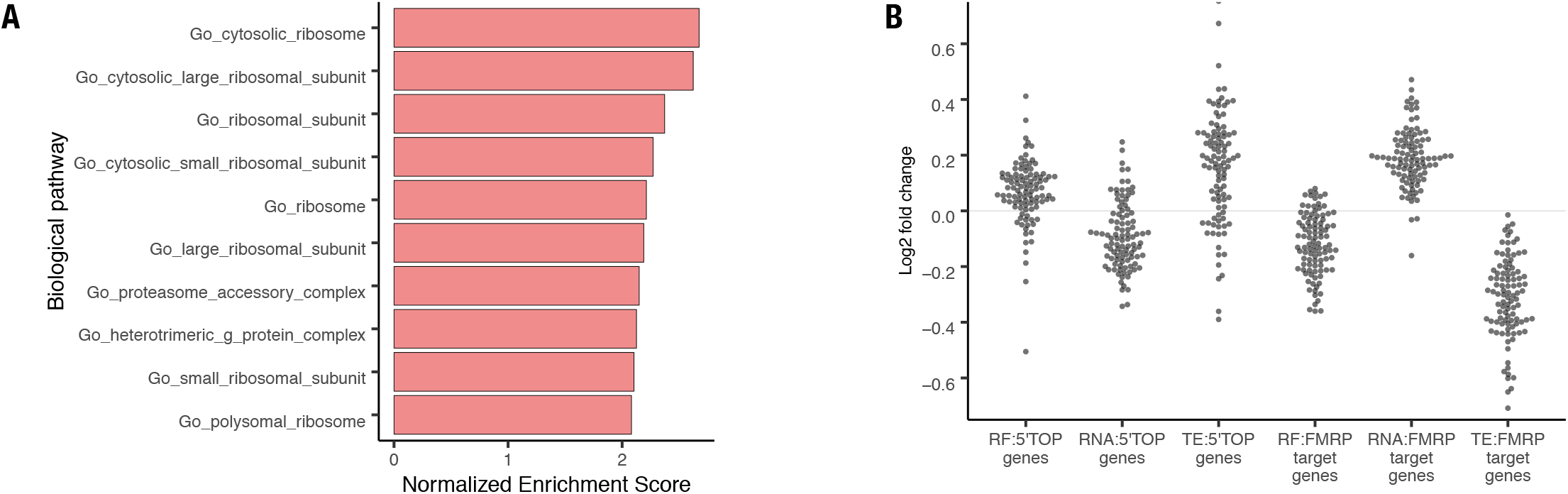
FXS brains exhibit differential alterations in translation of mRNAs with 5’ terminal oligopyrimidine motifs compared to mRNAs that are FMRP-binding partners. **(A)** The top ten gene ontologies exhibiting increased RF abundance in FXS cortices, when including all robustly measured genes in the analysis (instead of only using significantly different genes). mRNAs coding for ribosomal proteins, which are enriched in 5’TOP motifs, show increased RF abundance in FXS brains. **(B)** The log-fold-changes in RF abundance, mRNA expression and translation efficiency (TE) of the top 100 brain expressed 5’TOP motif bearing mRNAs, and the top 100 FMRP binding partner mRNAs, in FXS brains. The vast majority of mRNAs harboring 5’ TOP motifs exhibit elevated RF abundance, reduced mRNA expression, and increased translation efficiency in FXS brains. In contrast, the overwhelming majority of FMRP-binding-partner mRNAs exhibit reduced RF abundance, elevated mRNA expression, and reduced translation efficiencies in FXS brains.

